# Extreme Y chromosome polymorphism corresponds to five male reproductive morphs

**DOI:** 10.1101/2020.08.19.258434

**Authors:** Benjamin A Sandkam, Pedro Almeida, Iulia Darolti, Benjamin Furman, Wouter van der Bijl, Jake Morris, Godfrey Bourne, Felix Breden, Judith E. Mank

**Author notes:** Corresponding Author: Benjamin A Sandkam. Author Contributions B.A.S. and J.E.M. designed research; B.A.S., J.E.M, F.B., G.R.B. conducted field work; B.A.S., P.A., I.A., B.L.S.F., W.v.d.B., and J.M. conducted bioinformatic analyses; B.A.S., P.A., I.A., B.L.S.F., W.v.d.B., J.M., G.R.B., F.B. and J.E.M. wrote the paper.

## Abstract

Sex chromosomes form once recombination is halted between the X and Y chromosomes. This loss of recombination quickly depletes Y chromosomes of functional content and genetic variation, which is thought to severely limit their potential to generate adaptive diversity. We examined Y diversity in *Poecilia parae*, where males occur as one of five discrete morphs, all of which shoal together in natural populations where morph frequency has been stable for over 50 years. Each morph utilizes different complex reproductive strategies, and differ dramatically from each other in color, body size, and mating behavior. Remarkably, morph phenotype is passed perfectly from father to son, indicating there are five Y haplotypes segregating in the species, each of which encodes the complex male morph characteristics. Using linked-read sequencing on multiple *P. parae* females and males of all five morphs from natural populations, we found that the genetic architecture of the male morphs evolved on the Y chromosome long after recombination suppression had occurred with the X. Comparing Y chromosomes between each of the morphs revealed that although the Ys of the three minor morphs that differ predominantly in color are highly similar, there are substantial amounts of unique genetic material and divergence between the Ys of the three major morphs that differ in reproductive strategy, body size and mating behavior. Taken together, our results reveal the extraordinary ability of evolution to overcome the constraints of recombination loss to generate extreme diversity resulting in five discrete Y chromosomes that control complex reproductive strategies.

**Significance Statement:** The loss of recombination on the Y chromosome is thought to limit the adaptive potential of this unique genomic region. Despite this, we describe an extraordinary case of Y chromosome adaptation in *Poecilia parae*. This species contains five co-occurring male morphs, all of which are Y-linked, and which differ in reproductive strategy, body size, coloration, and mating behavior. The five Y-linked male morphs of *P. parae* evolved after recombination was halted on the Y, resulting in five unique Y chromosomes within one species. Our results reveal the surprising magnitude to which non-recombining regions can generate adaptive diversity and have important implications for the evolution of sex chromosomes and the genetic control of sex-linked diversity.

## Introduction

Sex chromosomes form when recombination is halted between the X and Y chromosomes. The loss of recombination results in a host of evolutionary processes that quickly deplete Y chromosomes of functional content and genetic variation, severely limiting the scope for adaptive evolution^1^. Y chromosomes can counter this loss to some degree through a variety of mechanisms^2-5^, however the adaptive potential of Y chromosomes is generally thought to be much lower than the remainder of the genome. Typically, this results in relatively low levels of Y chromosome diversity within species. The adaptive potential of non-recombining regions has far broader implications beyond just Y chromosomes given the increasing realization that supergenes, linked regions containing alleles at multiple loci underlying complex phenotypes, are key to many adaptive traits^6-12^. Many supergenes are lethal when homozygous and therefore also non-recombining^8,10^. Therefore, the processes that constrain Y chromosome evolution also affect much broader areas of the genome.

*Poecilia parae* is a small freshwater fish found in coastal streams of South America. Remarkably, males of this species are one of five distinct morphs that utilize different reproductive tactics, and differ in reproductive strategy, body size, color, and mating behaviour^13-19^ (summarized in Table S1). There are three major morphs: parae, immaculata and melanzona. The parae morph has the largest body, vertical black stripes, an orange tail-stripe and is highly aggressive, chasing away rival males and aggressively copulating with females by force. Immaculata, resembles a juvenile female. Although immaculata has the smallest body size of all morphs, it has the largest relative testes and produces the most sperm, employing a sneaker copulation strategy. Melanzona is sub-divided into three minor morphs, which are similar in body size and all have a colored horizontal stripe (either red, yellow or blue), which they present to females during courtship displays.

All five morphs co-occur in the same populations, and the relative frequency of morphs is highly stable over repeated surveys spanning 50 years (∼150 generations)^13,14,20^. This suggests that balancing selection, likely resulting from a combination of sexual and natural selection, is acting to maintain these five adapted morphs. Most importantly, multigeneration pedigrees show that morph phenotype is always passed perfectly from father to son^13^, indicating morphs are Y-linked. Given their discrete nature and complex phenotypes, it is clear that the five *P. parae* morphs are controlled by five different Y chromosomes. This system therefore offers the potential for a unique insight into the adaptive potential of Y chromosomes, and the role of these regions of the genome in male phenotypes.

We have recently shown that poecilid species closely related to *P. parae* share the same sex chromosome system as *Poecilia reticulata*^21^ (guppies), however the extent of Y chromosome degeneration differs markedly across the clade. Although the Y chromosome in *P. reticulata* and *Poecilia wingei* contains only a small area of limited degeneration^21-24^, the entirety of the Y chromosome of *Poecilia picta* is highly degenerate^21^. *P. parae* is a sister species of *P. picta*, however *P. picta* males are markedly different from *P. parae* and do not resemble any of the five *P. parae* morphs^*18,20,25-27*^, suggesting extreme diversity was generated on the *P. parae* Y chromosome after recombination was halted with the X chromosome. Work on model systems has indeed shown Y chromosomes can accumulate new genetic material^2-5^, yet these differences occur over long periods of time and are only evident when comparing Ys across species. Non-model systems, such as *P. parae*, provide a unique opportunity to explore the limits and role of non-recombining regions in generating diversity.

Because *P. parae* is very difficult to breed in the lab we collected tissue from natural populations in South America where all five male morphs co-occur and used linked-read sequencing on multiple females and males of all five morphs. We first confirmed that *P. parae* shares the same sex chromosome system as its close relatives^21,22^. We went on to find patterns of X-Y divergence are the same for all five Y chromosomes and matches the X-Y divergence we observed in *P. picta*, suggesting that the morphs emerged after Y chromosome recombination was stopped in a common ancestor of *P. parae* and *P. picta*. Comparing the five Y chromosomes to each other, we find that while the Ys of the three minor morphs (red, yellow and blue melanzona) that differ only in color are highly similar, the Ys of the three major morphs (parae, immaculata, and melanzona) that differ in reproductive strategy, body size and mating behavior are significantly diverged from one another and carry substantial amounts of unique genetic material. Taken together, our results reveal the surprising ability of the Y chromosome to not only overcome the constraints of recombination loss, but to generate extreme diversity, resulting in five discrete Y chromosomes that control complex reproductive strategies.

## Results

We collected 40 individual *P. parae* from natural populations in Guyana in December 2016, including eight red melanzona, four blue melanzona, five yellow melanzona, five immaculata, seven parae morph males, and 11 females. 29 samples with sufficiently high molecular weight were individually sequenced with 10X Genomics linked-reads. We generated a *de novo* genome assembly for each of these samples. The remaining 11 lower molecular weight samples were individually sequenced with Illumina sequencing paired end reads (see Tables S2 and S3 for sequencing and assembly details).

### The P. parae Y chromosome is highly diverged from the X and shared with P. picta

Degeneration of the Y chromosome results in reduced male coverage when mapped to a female reference genome. The ratio of male to female mapped reads can be used to identify regions where the Y and X chromosomes differ substantially from each other^21,28-30^. To do this, we used our best female *de novo P. parae* genome, based on N50 and other assembly statistics (see Table S3). We then determined chromosomal position of the scaffolds using the reference-assisted chromosome assembly (RACA) pipeline, which combines phylogenetic and sequencing data to place scaffolds along chromosomes^31^. Next we mapped reads from all 40 samples to this female assembly and calculated male:female coverage, first for each of the five morphs independently, and then all morphs together.

As we previously found in *P. picta*^*21*^, chromosome 8 (syntenic to *P. reticulata* chromosome 12) showed a clear signal of reduced read coverage in males (Figure 1a-b), indicating an XY sex determination system. Y divergence is evident across nearly the entire chromosome and is largely identical to the pattern we previously observed in *P. picta* (Figure 1c and Fig. S1). This suggests that these species inherited a highly degenerate Y chromosome from their common ancestor, well before the origin of the *P. parae* male morphs.

**Figure 1.**
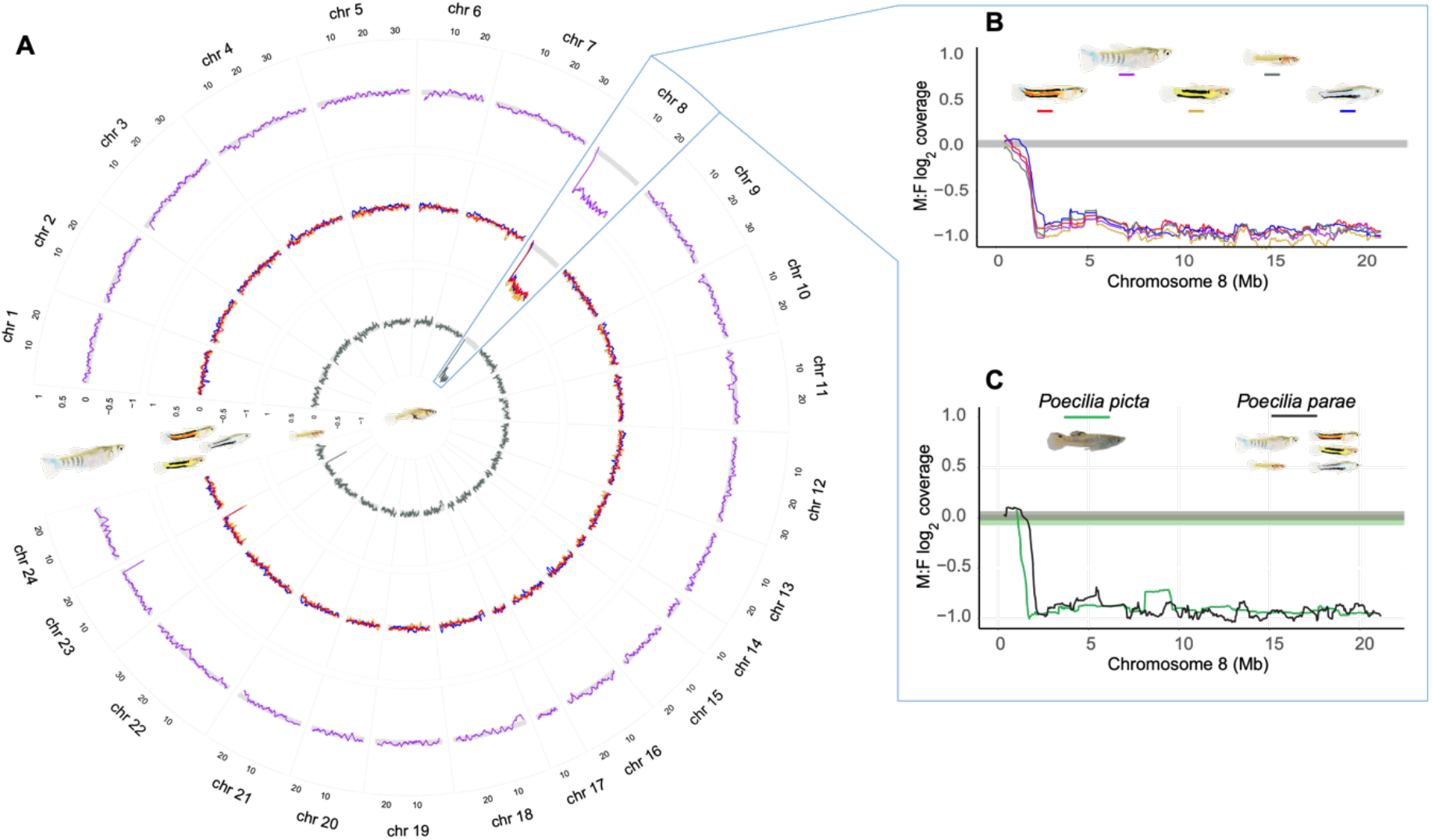
Coverage differences between the sexes (male:female log_2_) for female scaffolds placed by RACA on the reference *Xiphophorus hellerii* chromosomes. (A) Average immaculata (inner ring), the three melanzona (middle ring) and parae morphs (outer ring) plotted across all chromosomes. Highlighted in blue is *X. hellerii* chromosome 8 which is syntenic to the guppy sex chromosome (*P. reticulata* chromosome 12). The decreased male coverage of chromosome 8 indicates this is also the sex chromosome in *P. parae*. (B) All five *P. parae* morphs share the same pattern of XY divergence, indicating a shared history of recombination suppression. (C) Pattern of *P. parae* XY divergence is the same as the sister species *P. picta*, indicating recombination was stopped in the common ancestor of *P. parae* and *P. picta* (14.8-18.5 mya^43^). In each, horizontal grey-shaded areas represent the 95% confidence intervals based on bootstrap estimates across the autosomes.

Short sequences representing all the possible substrings of length *k* that are contained in a genome are referred to as *k-*mers, and *k*-mer comparisons between male and female genomes has been used to identify Y chromosome sequence (Y-mers) in a wide range of organisms^32-34^, including guppies^35^ and *P. picta*^21^. We first compared all males, representing all five morphs, to all females. We found a total of 27,950,090 Y-mers (of 31bp) that were present in at least two males but absent from all females. However, only 59 of these Y-mers were present in all 23 males (Figure 2). We found only 251,472 *k*-mers present in at least two females but absent from all males, and 0 of these were found in all 6 females, demonstrating our Y-mer approach had a very low false positive rate.

**Figure 2.**
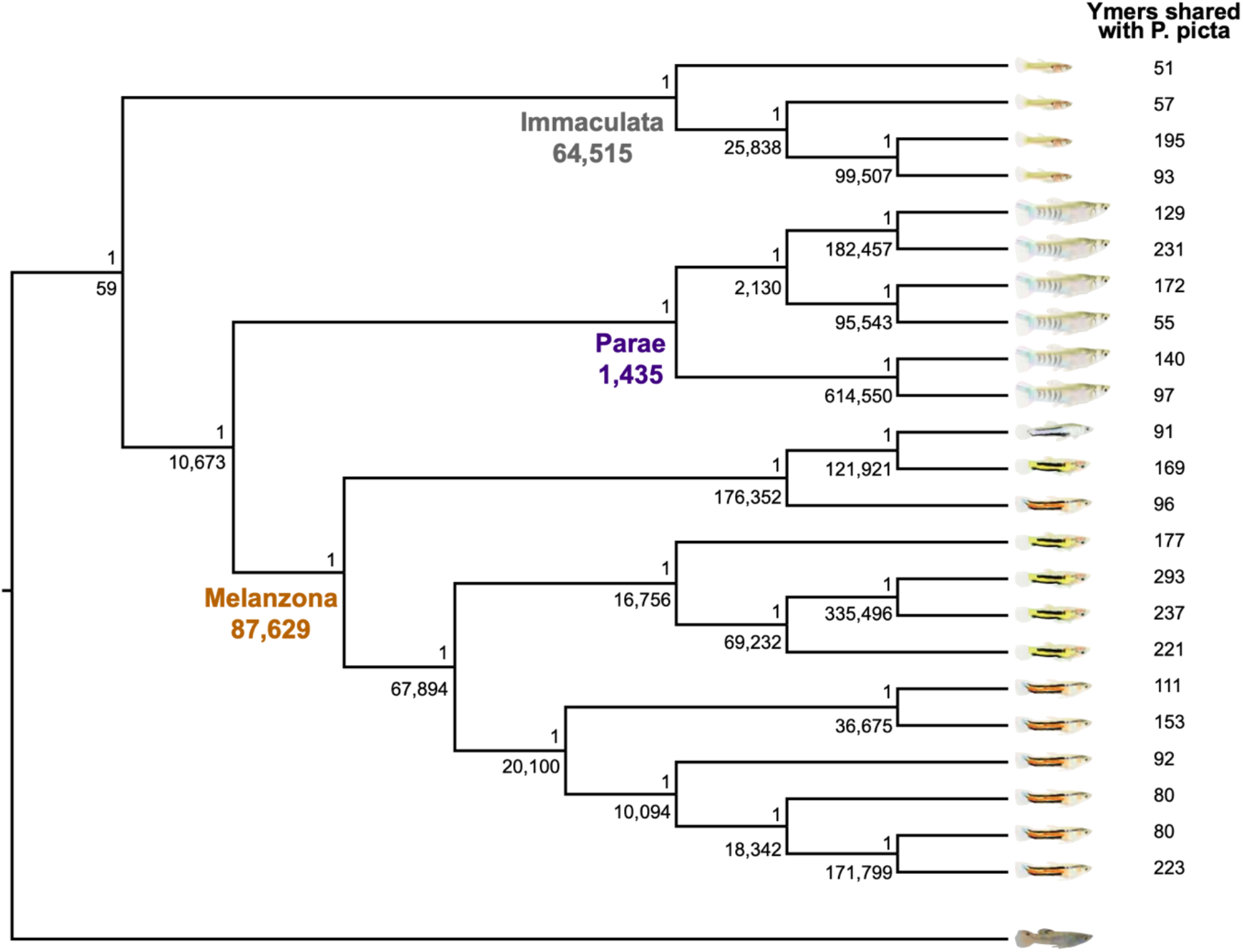
Bayesian Y chromosome phylogeny based on presence/absence of the 27,950,090 *P. parae* Y-mers and 1,646 *P. picta* Y-mers^21^ in each individual and rooted on *P. picta*. The posterior probability is presented above each node, below the node is the number of *P. parae* Y-mers unique to all members of that clade. The three major morphs of *P. parae* (immaculata, parae and melanzona) formed distinct clades and the Y-mers unique to all members of these clades are called morph-mers.

We next used Y-mer analysis to further test whether recombination was halted on the Y in the common ancestor of *P. parae* and *P. picta*. Of the 646,745 Y-mers that we previously identified in at least one *P. picta* male and no females^21^, 790 *P. picta* Y-mers matched Y-mers we identified in *P. parae*, consistent with a shared history of suppressed recombination. Additionally, these shared Y-mers were present in males of all morphs (Fig. S2) and discussed in more detail below. These shared Y-mers, combined with the striking similarity in male:female read mapping (Fig. S1) provide compelling evidence that the vast majority of Y chromosome recombination suppression occurred in a common ancestor of *P. picta* and *P. parae*.

### The P. parae Y chromosomes are highly diverged from each other

We next compared Y-mers across individuals, generating a phylogeny on the presence/absence of Y-mers in all the *P. parae* individuals with *P. picta* as an outgroup (Figure 2). Clear clades were recovered for each of the major morphs (immaculata, parae and melanzona) while the three minor morphs of melanzona (red, yellow, blue) were very similar to one another.

The phylogenic relationships of individuals (Figure 2) closely match the relative Y-mer comparisons across morphs (Figure 3). We found 64,515 Y-mers in every immaculata male that were not in any parae or melanzona males (i.e. immaculata-mers), 87,629 melanzona-mers, and 1,435 parae-mers, suggesting that the melanzona and immaculata Y chromosomes may contain more unique, non-repetitive sequence compared to the parae Y. Moreover, we found 10,673 Y-mers in all melanzona and parae males that did not occur in any immaculata males (Figure 2), suggesting that the parae Y shares greater sequence similarity to the melanzona Y.

**Figure 3.**
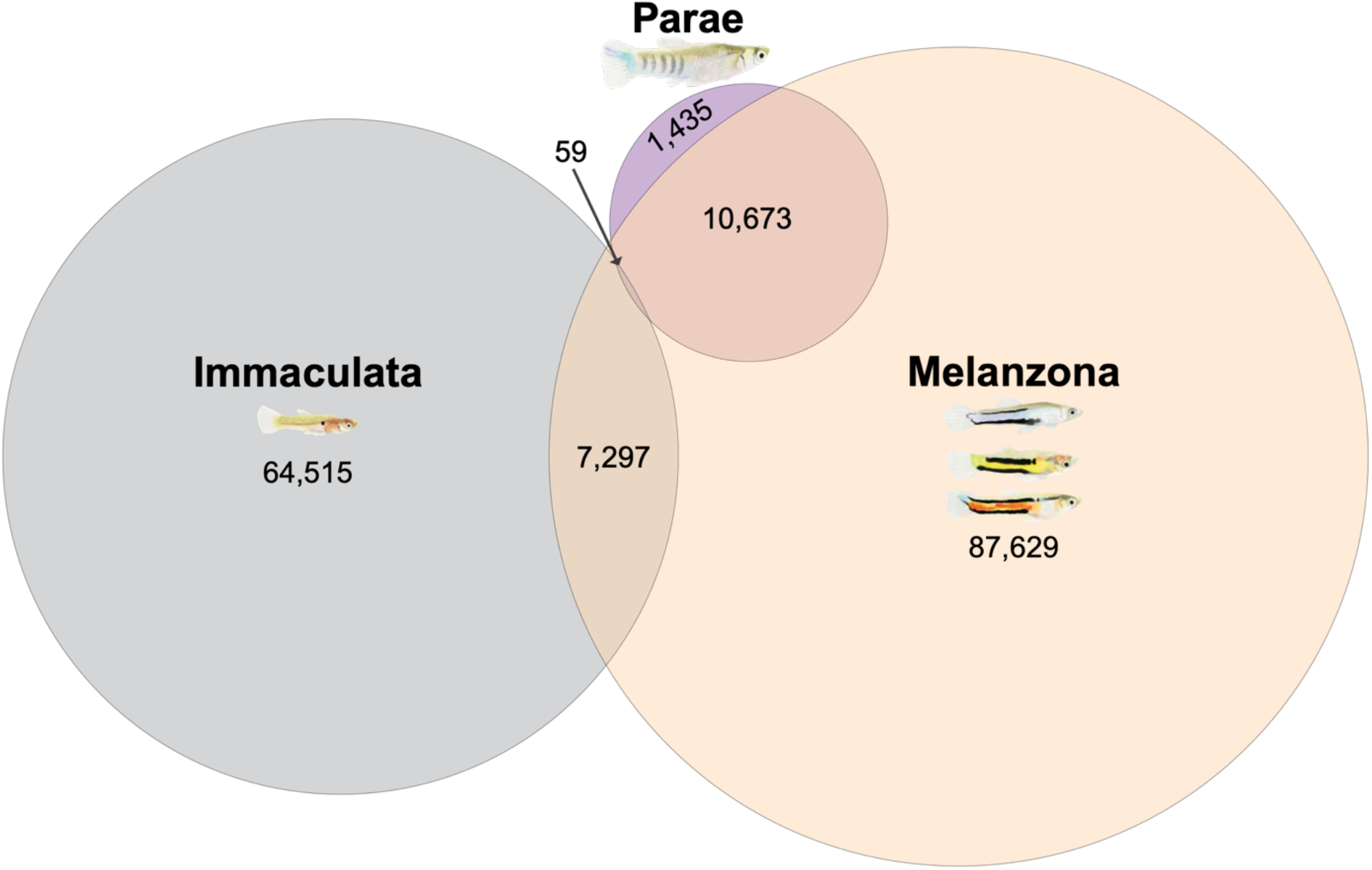
The distribution of the 27,950,090 *P. parae* Y-mers reveals strong differences across morphs. While there are very few Y-mers present in all morphs, each morph harbors unique Y-mers. The melanzona and parae morphs share more Y-mers with one another than either share with immaculata.

We calculated our false positive rate by randomly permuting our male samples into groups regardless of morph and determining Y-mers present in all males of each group that were absent from all other males. We found no unique Y-mers in groups of five or more random males, and just 31 unique Y-mers in groups of four random males, demonstrating the false positive rate of our morph-mer approach is exceedingly low (Fig. S3).

### Mapping morph-mers confirms high diversity of Y chromosomes

The large number of morph-mers we identified could either indicate that the discrete morphs are the result of low divergence across large Y chromosomes, or smaller complexes of highly diverged Y sequence. To resolve this, we mapped the respective set of morph-mers to the 21 *de novo* male genomes. If divergence across the morph-specific Y chromosomes is low compared to each other, mapped morph-mers would be dispersed across many scaffolds. Instead we found morph-mers were not evenly dispersed. For example, a single melanzona scaffold (∼110kb) containing 27% of all melanzona-mers (23,773), and most morph-mers overlapped one another (Fig. S4 and S5). This confirms our morph-mer approach identified complexes of highly diverged Y sequence.

To compare the relative size of these diverged complexes across morphs, we identified all scaffolds that contained >5 morph-mers in each individual. The average amount of sequence contained within these morph-mer scaffolds was 1.3 Mb for melanzona, 3.2 Mb for immaculata, and just 0.1 Mb for parae individuals (Table S4). This complements the relative number of Y-mers we found for each morph and together suggests the amount of unique Y chromosome sequence differs across morphs, with the parae morph Y containing the smallest amount of unique genetic material.

### Read mapping confirms high divergence of Y chromosomes

To determine how divergent the five Y chromosomes are from one another, we mapped reads from all 39 samples to full *de novo* genome assemblies of each male morph (200 total alignments). Most scaffolds contain autosomal sequence, and coverage is not expected to differ by sex or morph. Meanwhile, scaffolds containing morph specific sequence will have higher coverage by males of the same morph (e.g. immaculata reads mapped to an immaculata assembly) than coverage by males of a different morph (e.g. melanzona reads mapped to an immaculata assembly). Low female coverage of such scaffolds confirms these regions are on the Y and are substantially diverged from the X.

As expected, when comparing coverage between males of the same morph as the reference assembly and the other morphs we found average coverage was 1:1 when considering all scaffolds, yet scaffolds enriched for morph-mers (containing >5) had much higher coverage b males of the same morph as the reference (Figure 4). Surprisingly, we found no coverage by immaculata or parae reads for nearly half (40/100) of the scaffolds enriched for melanzona-mers. Similarly, 14 of the 93 scaffolds enriched for immaculata-mers had no coverage when we mapped melanzona and parae reads. Meanwhile, in agreement with our morph-mer analysis, all 12 of the scaffolds enriched for parae-mers had nearly equal coverage by melanzona and immaculata reads, once again suggesting that the parae Y contains very little unique Y sequence.

**Figure 4.**
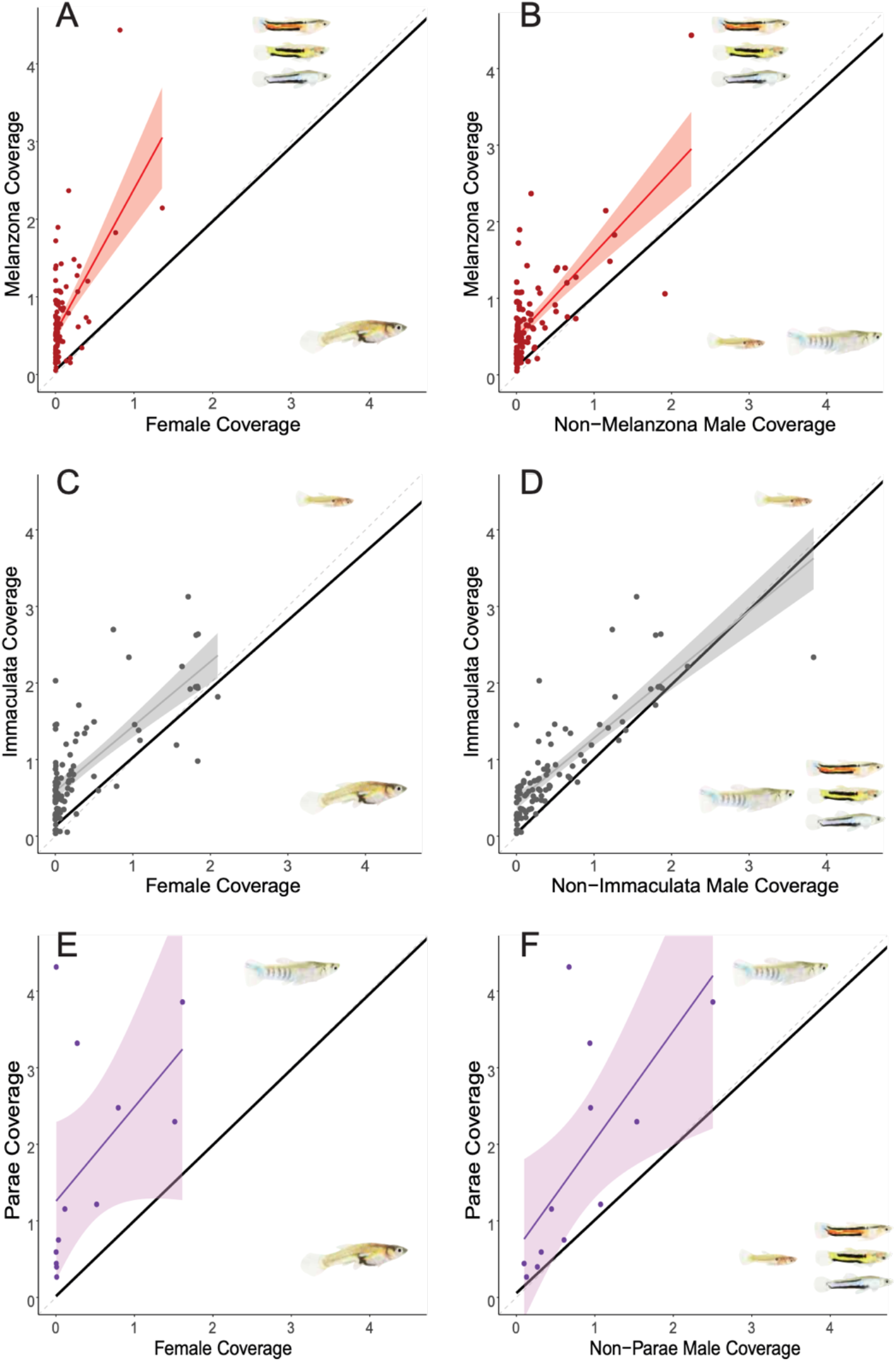
Relative corrected scaffold coverage of 39 individuals when aligned to melanzona (A-B), immaculata (C-D), and parae (E-F) *de novo* genomes. Scaffolds containing morph-mers had higher coverage by males than females (A,C,E) confirming these scaffolds contain male specific sequence. Scaffolds containing morph-mers also had higher coverage by males of the reference morph than males of the other morphs (B,D,F), indicating the Y chromosome sequence is substantially diverged across morphs. In each, corrected scaffold coverage of focal morph is on the Y axis and corrected scaffold coverage of the compared morph is on the X axis. The 1:1 line is denoted as a grey dashed line. The linear regression and standard error across all scaffolds are shown as a thick black line that is nearly 1:1 for all morphs (note – 95% confidence interval is presented but too small to distinguish from the regression line). The scaffolds containing morph-mers are shown as colored points, the linear regression and standard error of morph-mer scaffolds are shown as a colored line and shaded region respectively.

We also found the ratio of male:female coverage was much higher for morph-mer scaffolds, many of which had no female coverage, again confirming these complexes of morph specific sequence are located in non-recombining regions of the Y chromosome (Figure 4).

### Gene annotation of morph-mer scaffolds

We identified genes on scaffolds with >5 morph-mers. In total, we found 7 genes on the scaffolds containing the 59 Y-mers present in all morphs (totaling 30,558,901 bp), 291 genes on the immaculata scaffolds of sample P09 (totaling 9,748,162bp), 15 genes on the melanzona scaffolds of sample P01 (totaling 295,057 bp), and no genes on the parae morph scaffolds of P04 (totaling 127,542 bp) (Tables S5, S6 and S7).

Only one gene was predicted on scaffolds that were completely unique to melanzona (*trim35)*, and only two genes were predicted on scaffolds completely unique to immaculata (*trim39* and *nlrc3*). Members of the *Trim* gene family act throughout the body and are well known to rapidly evolve novel functions^36,37^. Meanwhile, *nlrc3* has been shown to selectively block cellular proliferation and protein synthesis by inhibiting the mTOR signalling pathway^38^, which could play a role in keeping immaculata the smallest morph. Several copies of the transcription factor *Tbx3* were present on male unique scaffolds in both melanzona and immaculata morphs. *Tbx* genes play key roles in development and act as developmental switches^39-41^, raising the possibility that it could play a role in orchestrating the multi-tissue traits that differ across morphs.

We also found several copies of *texim* genes on male scaffolds of melanzona and immaculata that most closely match *texim2* and *texim3*. While the function of *texim2* and *texim3* are largely unknown they have been shown to be highly expressed in the brain and testis of closely related species^42^. In *Xiphophorus maculatus*, a close relative to *P. parae* (∼45 mya^43^), the transposable element *helitron* has moved *texim1* to the sex determining region of the Y chromosome and duplicated it resulting in three copies of *texim* that are expressed specifically in late stage spermatogenesis^42^. The copies of *texim2* and *texim3* that we identified on the Y chromosome of *P. parae* are not the same as those in *X. maculatus* as the Ys arose independently and the *X. maculatus* Y is not chromosome 8.

We also found a large number of transposable elements on scaffolds enriched for the Y-mers that were present in all individuals, and the scaffolds enriched for morph-mers, including 90 copies of the *helitron* transposable element on scaffolds enriched for melanzona-mers, and 38 copies on scaffolds enriched for immaculata-mers (Tables S9 and S10). It is possible that a process similar to *X. maculatus* has occurred in *P. parae* where either TEs have moved *texim2* and *texim3* to Y specific sequence or Y specific sequence has evolved around these genes. Future analyses are needed to determine the roles of these and other genes in generating the morph-specific phenotypes.

## Discussion

Recombination is widely regarded as one of the most important processes generating phenotypic diversity as it produces novel allelic combinations on which selection can act^44^. The loss of recombination is classically assumed to prevent the generation of novel large-scale phenotypes, as non-recombining regions are expected to rapidly lose diversity through sweeps and background selection^1,45-53^ and have only limited potential for adaptive evolution. The power of these processes to deplete non-recombining regions of diversity is clearly evident in the Y chromosomes of many species^1^.

Our results, based on both similarity of Y degeneration (Fig. S1) and shared Y-mers, are consistent with recombination suppression between the X and Y chromosome in the common ancestor of *P. parae* and *P. picta* (14.8-18.5 mya^43^) and a highly degenerate Y present at the origin of *P. parae*. Importantly, none of the characteristics that differentiate the immaculata, parae or three melanzona morphs of *P. parae* are found in any close relatives, which means the genetic basis of the extreme diversity in morphs evolved on a non-recombining, highly degenerate Y chromosome. Given that these morphs differ in a suite of complex traits, including body size, testis size, color pattern, and mating strategy, it is highly likely that they are underpinned by polygenic genetic architectures, which evolved on the *P. parae* Y chromosome. Consistent with complex differences between morphs, we identified substantial morph-specific genetic material (Figures 2-4) that was also absent in females and therefore Y-linked.

Although male-specific regions of the genome may experience elevated mutation rates^54^, it is far more likely that the high *P. parae* Y diversity was generated through translocations and/or transposable element (TE) movement. Translocations have been shown to increase Y-chromosome content^2,55^. In contrast, although TE movement has historically been considered to be a deleterious process, more recent reports have revealed that TEs can alter regions by removing or adding regulatory or coding sequence^39,56-59^, and TEs may even act as substrate for novel genes^60^. As predicted for non-recombining regions, we found a large number of TEs in the five *P. parae* Y chromosomes.

### Making and Maintaining Five Morphs

We found remarkable morph specific genetic diversity on the Y chromosome of *P. parae*. Intriguingly, that diversity is maintained within populations, as evidenced by the stability in morph frequencies over repeated surveys spanning 50 years, or roughly 150 generations^13,14,20^. Even if alternative morphs have exactly equal fitness, populations are expected to eventually fix for one morph due to drift^61^. Maintaining alternative morphs within the same population relies on negative frequency dependent selection, thus as one morph decreases in frequency its fitness increases, such as with the three male morphs of the side blotched lizard^62^.

Previous work suggests that *P. parae* morphs are also under negative frequency dependent selection^14-17^, and this could facilitate the establishment and maintenance of five distinct Y chromosomes within the same species. Most new mutations are expected to be lost through drift if they do not confer a high enough fitness advantage over alternative alleles^63^, but mutations resulting in a new morph would be at the lowest frequencies and thus have the highest fitness, allowing them to rapidly stabilize in the population. Alternatively, it is possible that the morphs arose in separate populations and only later came into sympatry.

### Genetic Basis of Male Reproductive Morphs

Autosomal non-recombining regions have been shown to be associated with alternative reproductive strategies in a range of species^7,9,64-68^. For example, male morphs of the white throated sparrow, which differ in pigmentation and social behavior, are the result of a hybridization event which instantly brought together alternative sequence and halted recombination^65^. Importantly, because none of the characteristics of the *P. parae* morphs are found in any close relatives, it is unlikely that hybridization is the source of the Y chromosomes we describe.

The alternative male morphs in the ruff are controlled by an autosomal supergene that is composed of two alternative versions of an inverted region^10^. It has yet to be determined whether the diversity across these ruff supergenes pre-dates the inversion or arose after recombination stopped as it did in *P. parae*.

A large inverted region is also associated with social morphs in many ant species, this region formed in a common ancestor and has been maintained by balancing selection through repeated speciation events^12^. Although it is possible that the multiple male morphs in *P. parae* arose in an ancestor, this is less likely as they have not been observed in any related species to date.

## Conclusion

The role of recombination in shaping co-adapted allele complexes has long been an enigma, given that recombination is both a key mechanism in generating diverse allelic combinations, yet recombination also acts to break up such combinations. Here we found that tremendous diversity can still be generated without the power of recombination, and the Y chromosome contains remarkable adaptive potential with regard to male phenotypic evolution. The five Y-linked male morphs of *P. parae* emerged and diverged after recombination was halted, resulting in five unique Y chromosomes within one species. Future work identifying the mechanisms by which morphs are determined by these five Y chromosomes will provide much needed insight to determining which evolutionary forces have led to and shaped these amazing complexes and their co-evolution with the rest of the genome, which is shared across all morphs.

## Methods

### Field Collections and DNA Isolation

To ensure we accounted for natural diversity in the five Y chromosomes of *P. parae*, and because this species is extremely difficult to breed in captivity, we collected all samples (N=40) from large native populations around Georgetown, Guyana in 2016 (see Table S2 and Sandkam, et al. ^18^ for description of populations) (Environmental Protection Agency of Guyana Permit 120616 SP: 015). Individuals were rapidly sacrificed in MS-222, whole-tail tissue was dissected into EtOH and immediately placed in liquid nitrogen to maintain integrity of high molecular weight (HMW) DNA. Tissue samples were brought back to the lab and kept at -80° C until HMW DNA extraction.

HMW DNA was extracted from 25mg tail tissue of each sample following a modified protocol from 10X Genomics described in Almeida *et al*^24^. Briefly, nuclei were isolated by gently homogenizing tissue with a pestle in cold Nuclei Isolation Buffer from a Nuclei PURE Prep Kit (Sigma). Nuclei were pelleted and supernatant removed before being digested by incubating in 70 µl PBS, 10 µl Proteinase K (Qiagen), and 70 µl Digestion buffer (20mM EDTA, 2mM Tris-HCL, 10mM N-Laurylsarcosine) for two hours at room temperature on a tube rotator. Tween 20 was added (0.1% final concentration) and DNA was bound to SPRIselect magnetic beads (Beckman Coulter) for 20 min. Beads were bound to a magnetic rack and washed twice with 70% Ethanol before eluting DNA. Samples were visually screened for integrity of HMW DNA on an agarose gel. Of the 40 samples extracted, 29 passed initial screening and were used for individual 10X Chromium linked-read sequencing (six female, seven red, five yellow, one blue, four immaculata, and six parae morph). The remaining 11 samples (five female, one red, three blue, one immaculata, and one parae morph) were individually sequenced on an Illumina HiSeqX as 2 x 150 bp reads using the v2.5 sequencing chemistry with 300bp inserts and trimmed with trimmomatic (v0.36)^69^. To ensure high coverage of the Y, all 40 samples were individually sequenced to a predicted coverage of 40X (see Table S2 for number of reads after filtering), which would result in predicted 20X coverage of the haploid Y.

### 10X Chromium Linked-read Sequencing and Assembly

10X Chromium linked-read sequencing was performed at the SciLifeLab, Uppsala Sweden. The 10X Chromium pipeline adds unique tags to each piece of HMW DNA before sequencing on an Illumina platform. These tagged reads were either used directly in the 10X assembly pipeline, or tags were removed using the *basic* function of Longranger v.2.2.2 (10X Genomics), trimmed with trimmomatic (v0.36)^69^ and treated as normal Illumina reads^70^ for coverage analyses (Table S2).

Scaffold level *de novo* genomes were assembled for each of the 29 linked-read samples using the Supernova v2.1.1 software package (10X Genomics) (see Table S3 for assembly statistics).

A female chromosome level assembly was created by assigning scaffolds to chromosomal positions. The female with the best *de novo* assembly (largest assembly size and scaffold N50) was used for the Reference Assisted Chromosome Assembly (RACA) pipeline^31^. Briefly, scaffolds were aligned with LASTZ v1.04^71^ against high-quality chromosome level genome assemblies of a close relative (*Xiphophorus helleri* v4.0; GenBank accession GCA_003331165) and an outgroup (*Oryzias latipes* v1; GenBank accession GCA_002234675). Alignments were then run through the UCSC chains and nets pipeline from the kentUtils software suite^72^ before passing to the RACA pipeline. RACA uses alignments of short-insert and long-insert paired reads that bridge scaffolds to further order and confirm scaffold arrangement. For short-insert data, 150bp reads from the five females sequenced with paired-end Illumina (300bp inserts) were aligned to the target assembly with Bowtie2 v2.2.9^73^ reporting concordant mappings only (--no-discordant option). For long-insert data, synthetic 150bp ‘pseudo-mate-pair’ reads were generated from the *de novo* scaffolds of the 6 female *P. parae* samples sequenced with Chromium linked-reads. To increase the likelihood that bridge reads spanned scaffolds, we generated two long-insert pseudo-mate-pair libraries for each of the six *de novo* female genomes, a 2.3kb insert library and a 15kb insert library, and aligned these to the target assembly. RACA then used the information from both the phylogenetically weighted genome pairwise alignments, and the read mapping data to order the target scaffolds into longer predicted chromosome fragments.

To identify which chromosome is the sex chromosome and determine the extent of X-Y divergence, we mapped reads from all 40 individuals to the female scaffolds that had RACA generated chromosome annotations using the *aln* function of *bwa* (v0.6.1)^74^. Alignments were filtered for uniquely mapped reads and average scaffold coverage was calculated using soap.coverage v2.7.7 (http://soap.genomics.org.cn/). To account for differences across individuals in sequencing library size, we divided the coverage of each scaffold by the average coverage across all scaffolds for each individual. Male to female (M:F) fold change in coverage of each scaffold was calculated for all males, and each of the five morphs as log_2_(average male coverage) – log_2_(average female coverage)^75^. Upon observing chromosome 8 was the sex chromosome, we calculated the 95% CI for M:F coverage by bootstrapping across all scaffolds which RACA placed on the autosomes.

### Identifying Morph specific sequence by k-mers

To locate morph specific sequence, we identified morph specific *k*-mers (morph-mers) and then mapped these to the respective *de novo* genome assemblies. First Jellyfish v2.2.3^74^ was used to identify all 31bp *k*-mers from the ‘megabubble’ output of each of the 22 male *de novo* genome assemblies from supernova. The ‘megabubble’ outputs from supernova include all sequence before being flattened into final phased scaffolds, this minimizes any potential *k*-mer loss due to arbitrary flattening (a benefit over *k*-mers from the full psuedohap output of supernova) and excludes *k*-mers from sequencing errors (a benefit over *k*-mers from sequencing reads). We next identified the putative Y-linked *k*-mers in each sample by removing all *k*-mers present in any of the females. For female *k*-mer identification we conservatively took *k*-mers from both the megabubble outputs and the raw Illumina reads of all 11 female individuals, which we used to identify every 31bp *k*-mer present >3 times (this maximized the chance of including *k*-mers from unassembled regions of female genomes but minimized *k*-mers from sequencing errors^35^). The *k*-mers present in females all occur either on autosomes or the X chromosome, therefore by removing the female *k*-mers from all *k*-mers identified in males we are left with putative Y-linked *k*-mers that we call Y-mers^21,35^. To exclude *k*-mers representing autosomal SNPs unique to a single male we required Y-mers be present in at least two individuals. Since all female sequence is theoretically present in males, we validated our method by identifying all *k*-mers from the six female megabubbles that were not present in the male Illumina reads and found how many were present in at least two female individuals.

We then combined all the Y-mers we found with those we previously identified in *P. picta*^21^ and used the presence of Y-mers as character states to build a phylogeny of all individuals and *P. picta*. Two runs of MrBayes v3.2.2^76^ were run for 100,000 generations with Y-mers treated akin to restriction sites (model F81 with rates set to equal). The SD of the split frequencies between runs reached 0 indicating both runs converged on identical and robust trees.

Monophyletic clades were recovered for each of the major morphs (immaculata, parae and melanzona). We then identified unique Y-mers in each clade (Y-mers present in every individual of that clade but not present outside that clade). This approach provided us with all morph-mers (Y-mers present in every individual of a given morph but not present in any individual of the other morphs).

These morph-mers reveal two insights: (1) at a gross level they provide a sense of how much Y sequence is shared within versus across morphs and (2) mapping these morph-mers to the respective *de novo* genomes allows us to identify regions of morph specific sequence^34,35,77,78^. To find these regions of morph specific sequence we first mapped the corresponding set of 31bp morph-mers to each of the 22 *de novo* male genomes (pseudohap style of Supernova output) using bowtie2^73^ allowing for no mismatches, gaps, or trimming. We found morph-mers disproportionately map to scaffolds, indicating they came from regions of highly diverged morph-specific sequence rather than evenly dispersed lowly diverged sequence (Fig. S3 and S4).

To verify our pipeline was identifying true Y-specific alignments we attempted to align the Y-mers and all of the morph-mers to each of the six female *de novo* genome assemblies and found no alignments could be made. We next verified that our morph-mers were targeting morph specific sequence by attempting to align each of the morph-mer datasets to individuals of the opposite morphs and again found no alignments could be made.

### Coverage analysis

To independently verify that the scaffolds identified by our *k*-mer approach contained highly diverged sequence, we aligned each of the 39 individuals to the individual with the best *de novo* genome of each morph (based on assembly size, scaffold N50, and contig N50, Supplementary Table 3). Alignments were generated with the *mem* function of *bwa* (v0.7.17)^74^. *samtools* (v1.10) was used to remove unmapped reads and secondary alignments with the *fixmate* function, and duplicates were removed with *markdup. bamqc* was then used to assess distribution of map quality. For each individual, the average coverage of each scaffold by reads with mapq >60 was determined using the *depth* function of *samtools* (v1.10). Y chromosomes are notorious for high incidents of transposable elements and repeats, this highly conservative filtering decreased false alignments to these regions. To account for differences across individuals in sequencing library size, we took the coverage of each scaffold divided by that individual’s coverage across all scaffolds. The average raw scaffold coverage across all individuals was 29.97X, therefore any scaffold with a corrected coverage < 0.025 (raw coverage <1X) was considered to have a coverage of 0.

### Gene Annotation

To identify genes on morph specific scaffolds we followed the pipeline described in Almeida *et al*^24^. Briefly, we took a very conservative approach by annotating only the scaffolds with >5 morph-mers from each of the *de novo* references used for the coverage analysis (one of each morph). The chance of a scaffold containing any particular 31bp *k*-mer depends on the length of the scaffold and can be calculated roughly as 0.25^31^ x scaffold length. The longest male scaffold we recovered was 19,887,348 and therefore had the greatest chance of containing any given Y-mer; 4.31*10^−12^. The most abundant morph-mers were melanzona-mers (87,629), therefore the likelihood of the largest scaffold containing one melanzona-mer by chance was 4.31*10^−12^ x 87,629 = 3.78*10^−7^ and the likelihood it contains 5 melanzona-mers purely by chance was roughly 7.71*10^−33^. If we conservatively assume that all scaffolds have the same probability of containing a Y-mer by chance and the male with the most scaffolds had 25,416 scaffolds – there was a likelihood of 1.96*10^−28^ that a scaffold was incorrectly identified.

We then annotated these scaffolds with MAKER v2.31.10^79^. We ran the MAKER pipeline twice: first based on a guppy-specific repeat library, protein sequence, EST and RNA sequence data (later used to train *ab-initio* software) and a second time combining evidence data from the first run and *ab-initio* predictions. We create a repeat library for these scaffolds using *de novo* repeats identified by RepeatModeler v1.0.10^80^ which we then combined with Actinopterygii-specific repeats to use with RepeatMasker v4.0.7^81^. Annotated protein sequences were downloaded from Ensembl (release 95)^82^ for 8 fish species: *Danio rerio* (GRCz11), *Gasterosteus aculeatus* (BROADS1), *Oryzias latipes* (ASM223467v1), *Poecilia latipinna* (1.0), *Poecilia mexicana* (1.0), *Poecilia reticulata* (1.0), *Takifugu rubripes* (FUGU5) and *Xiphophorus maculatus* (5.0). For ESTs, we used 10,664 tags isolated from guppy embryos and male testis^83^. Furthermore, to support gene predictions we also used two publicly available libraries of RNA-seq data collected from guppy male testis and male embryos^84^ and assembled with StringTie 1.3.3b^85^. As basis for the construction of gene models, we combined *ab-initio* predictions from Augustus v3.2.3^86^, trained via BUSCO v3.0.2^87^, and SNAP v2006-07-28^88^. To train Augustus and SNAP, we ran the MAKER pipeline a first time to create a profile using the protein and EST evidence along with RNA-seq data. Both Augustus and SNAP were then trained from this initial evidence-based annotation. Functional inference for genes and transcripts was performed using the translated CDS features of each coding transcript. Gene names and protein functions were retrieved using BLASTp to search the Uniprot/Swissprot, InterProscan v5 and GenBank databases.

## Data Availability

All reads will be submitted to GenBank.

## Acknowledgements

We thank the members of the Mank lab and Dr. Nora Prior for stimulating conversations and excellent feedback on early drafts of the manuscript. This was supported by the Natural Sciences and Engineering Research Council of Canada through a Banting Postdoctoral Fellowship (to B.A.S.), the European Research Council (Grants 260233 and 680951 to J.E.M.), and a Canada 150 Research Chair (to J.E.M.). Field work was conducted under Permit 120616 SP: 015 from the Environmental Protection Agency of Guyana. Sequencing was performed by the SNP&SEQ Technology Platform in Uppsala, Sweden. The CEIBA Biological Center partially subsidized our expenses during field collection in Guyana. We thank Clara Lacy for the illustrations of *P. parae*.

