## Supplemental Material for "Extreme Y chromosome polymorphism corresponds to five male reproductive morphs"

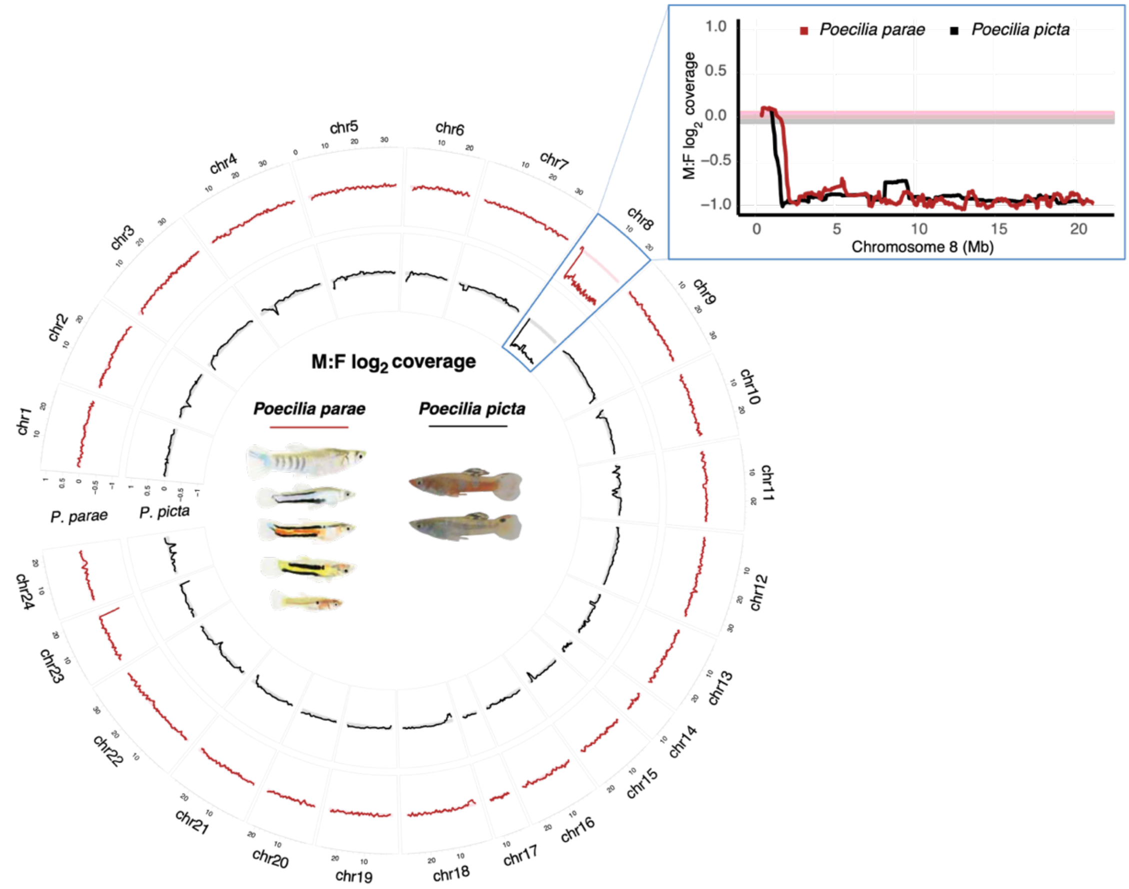


**Fig. S1.** Divergence between X and Y in *Poecilia parae* and the sister species *Poecilia picta* indicate recombination was stopped before the five morphs controlled by the Y chromosome evolved in *Poecilia parae*. M:F $\log_2$ coverage of RACA anchored scaffolds for all five morphs of *P. parae* (red) and the close relative *P. picta* (black) (Darolti *et al.* 2019). Lines represent sliding window of 15 scaffolds. Shaded bars represent the 95% confidence interval based on bootstrapping coverage across the autosomes for *P. parae* (pink) and *P. picta* (grey).

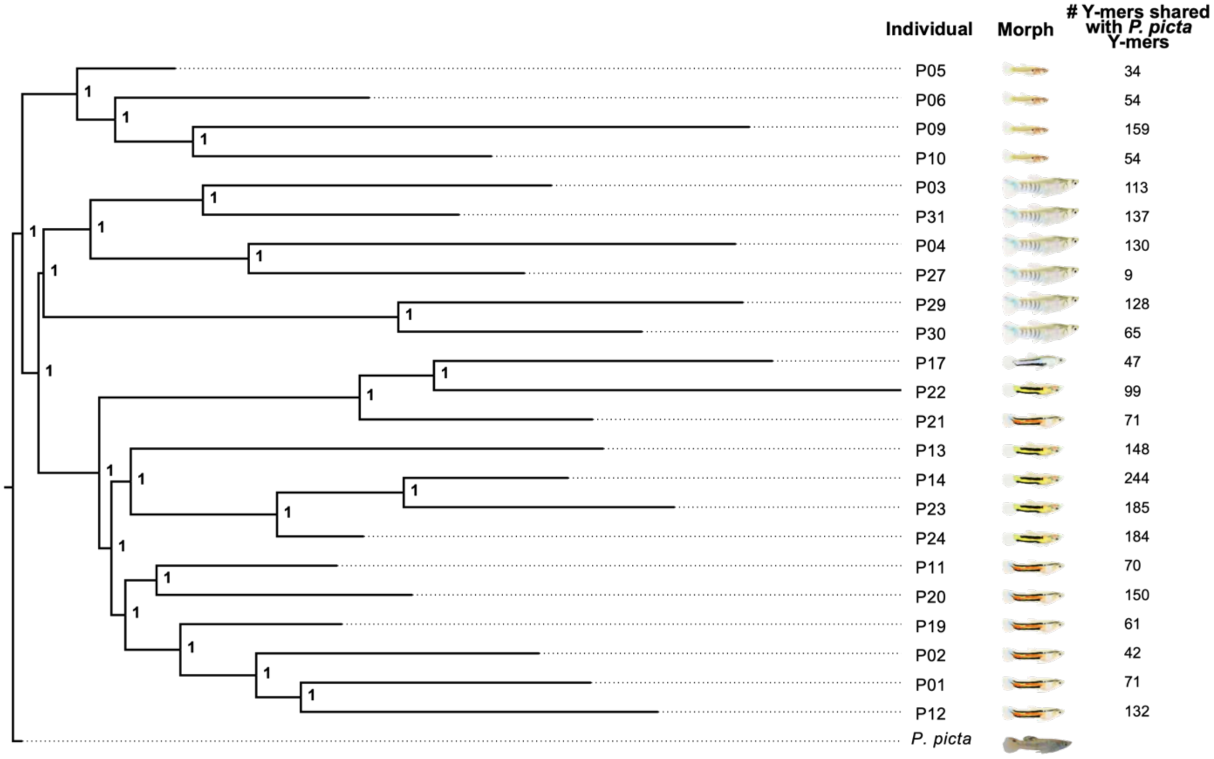


**Fig. S2.** Bayesian phylogeny built on presence/absence of the 27,950,090 *P. parae* Y-mers and the 646,754 *P. picta* Y-mers in each individual and rooted on *P. picta* (as in Figure 2). The posterior probability is presented at each node. The number of Y-mers each individual shares with *P. picta* Y-mers is denoted to the right. *P. picta* Y-mers are distributed across all morphs indicating that they have been segregating on non-recombining regions of the Y chromosome since recombination was stopped in the common ancestor of *P. parae* and *P. picta*.

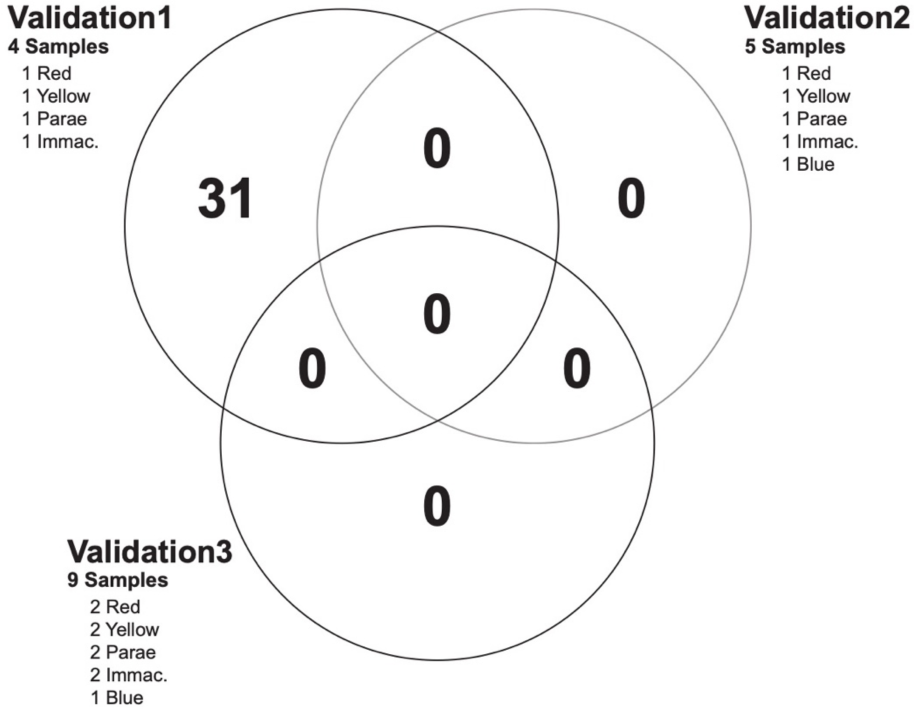


**Fig. S3.** Validation of morph-mer identification pipeline using random sets of individuals from each of the different morphs. Different samples were used for each set except for blue where the 1 sample was used in validation set 2 and validation set 3. There were a low number of Y-mers unique to sets of four random individuals and zero Y-mers unique to sets with more than four individuals. This demonstrates the false positive rate of our analysis was quite low because all major morphs had at least four individuals.

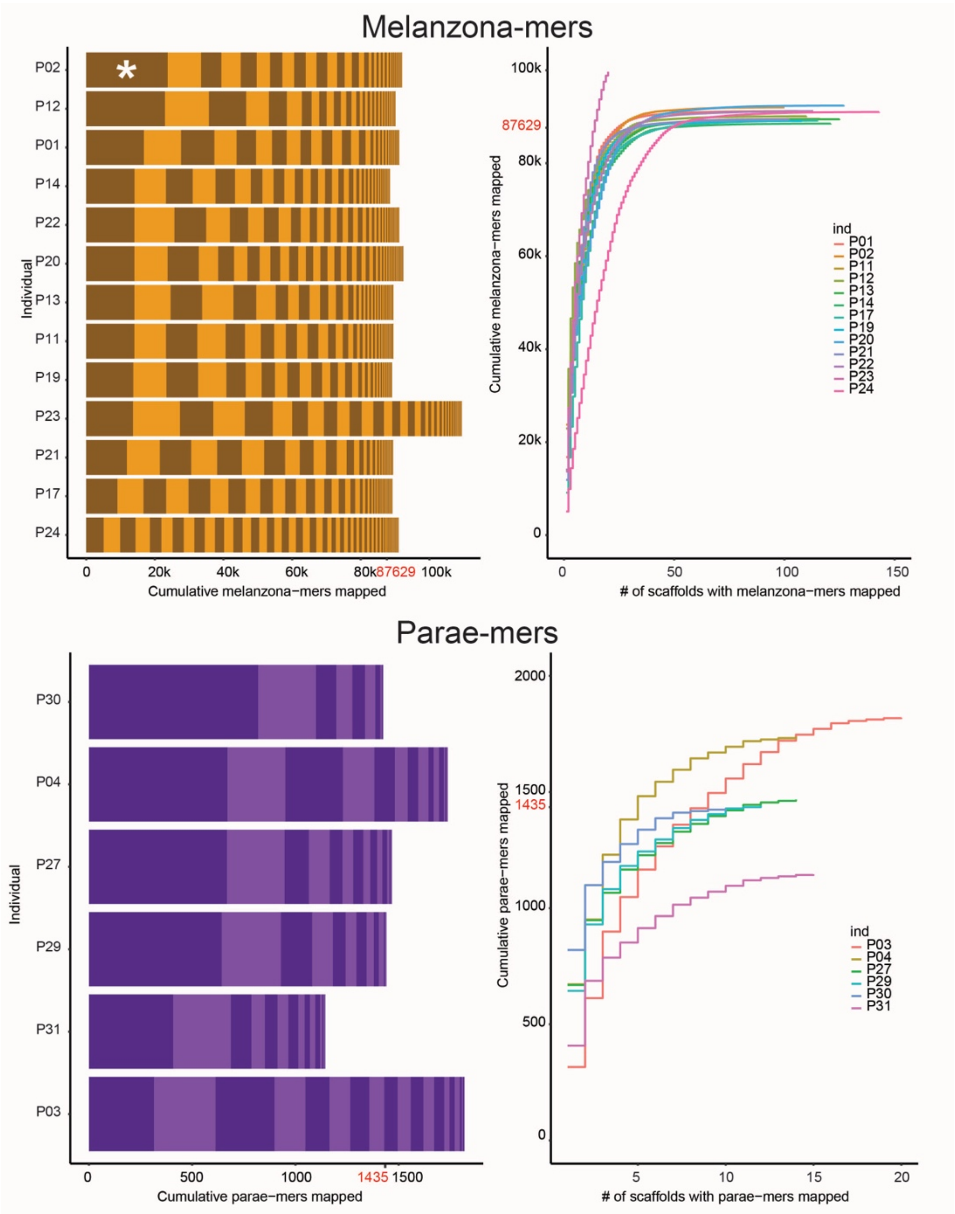

Melanzona-mers
P02
P12
P01
P14
P22
P20
P13
P11
P19
P23
P21
P17
P24
Individual
*
0
20k
40k
60k
80k
87629
100k
Cumulative melanzona−mers mapped
100k
87629
80k
60k
40k
20k
0
Cumulative melanzona−mers mapped
0
50
100
150
### of scaffolds with melanzona−mers mapped
ind
P01
P02
P11
P12
P13
P14
P17
P19
P20
P21
P22
P23
P24
Parae-mers
P30
P04
P27
P29
P31
P03
Individual
0
500
1000
1435
1500
Cumulative parae−mers mapped
2000
1500
1435
1000
500
0
Cumulative parae−mers mapped
5
10
15
20
### of scaffolds with parae−mers mapped
ind
P03
P04
P27
P29
P30
P31

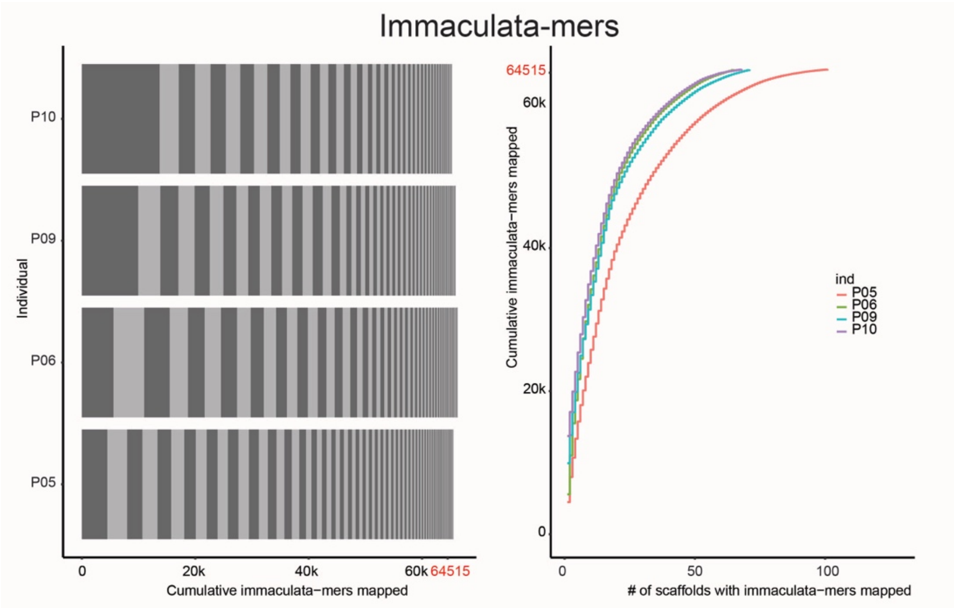


**Fig. S4.** Mapping distribution for each set of morph-mers mapped to *de novo* scaffolds of males of that morph with no mismatches, gaps, or trimming. Left: cumulative morph-mers mapped for each individual, each change in hue is a different scaffold. A large percentage of morph-mers generally map to just one or a few scaffolds indicating that our *k*-mer approach reveals regions of highly diverged morph specific sequence rather than single SNPs distributed throughout the genome. Right: cumulative morph-mers mapped presented as a function of the number of scaffolds. The strong deviation from 1:1 shows morph-mer mapping is non-random and further supports the morph-mers approach is identifying regions of morph specific sequence. The total number of unique morph-mers identified for that morph is indicated in red on the axis (note the variation in number of morph-mers mapped is due to some individuals having morph-mers map to multiple scaffolds). Astrix in P02 of the melanzona-mers indicates the example alignment scaffold with melanzona-mers presented in Supplementary Figure 4.

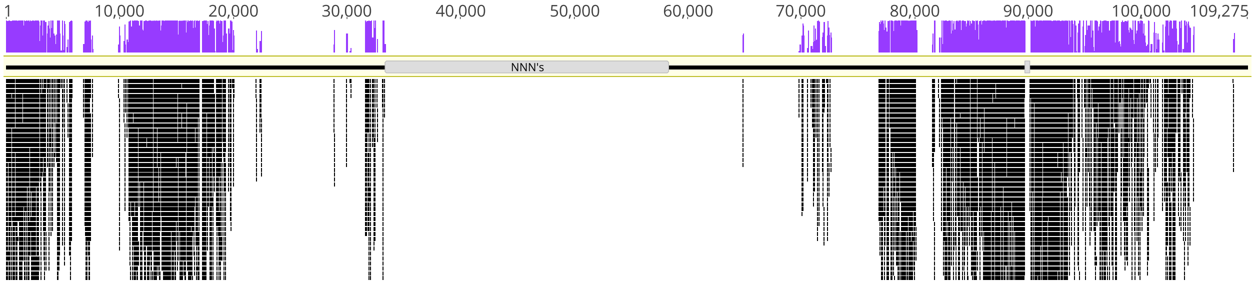


**Fig. S5.** Melanzona-mers aligned to scaffold 104666 of sample P02 with no mismatches, gaps, or trimming. Each 31bp melanzona-mer is shown aligned below the reference sequence, and coverage is shown in purple above the reference sequence. Of the 87,629 unique melanzona-mers; 23,773 aligned to this scaffold. Regions of Ns are denoted on the reference genome in grey and could explain a lack of melanzona-mers aligning to these regions. The strong clustering and overlapping nature of the melanzona-mers indicates sequence is highly diverged both from females and from the other morphs.

**Table S1.** Phenotypic characteristics that differ across the five male morphs of *Poecilia parae*.

| | Parae 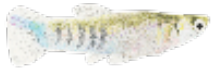 | Immaculata 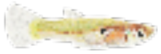 | Melanzona Red 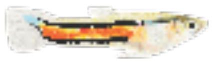 | Melanzona Yellow 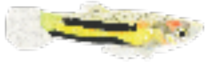 | Melanzona Blue 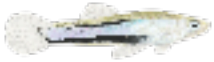 | Ref. |
|---|---|---|---|---|---|---|
| *Physiology* | | | | | | |
| **Body Size** | Large | Small | Medium | Medium | Medium | 1 |
| **Color Pattern** | Vertical stripes | Resembles juvenile female | Horizontal stripes | Horizontal stripes | Horizontal stripes | 1,2,3 |
| **Coloration** | Silver/purple | None | Red | Yellow | Blue | 1,2,3 |
| **Absolute Testis Mass** | Medium (90% of immaculata) | Largest | Small (70% of immaculata) | Medium (90% of immaculata) | Small (70% of immaculata) | 1 |
| **Testis/Body Mass** | Intermediate | High | Low | Intermediate | Low | 1 |
| **Sperm Midpiece** | Long | Short | Long | Long | Long | 1 |
| **Flagellum Length** | Short | Long | Short | Intermediate | Short | 1 |
| **Sperm per Ejaculate** | Fewer | Most | Fewer | Fewer | Fewer | 1 |
| **Sperm Length** | Short | Long | Short | Long | Short | 1 |
| *Behavior* | | | | | | |
| **Interaction w/ other males** | Highly aggressive | Non-aggressive | Aggressive | Aggressive | Non-aggressive | 3 |
| **Courtship Behavior** | Force copulate | Sneak copulate | Display | Display | Display | 1 |

(1) Hurtado-Gonzales and Uy. 2009. *Anim. Behav*. **77**,1187-1194. (2) Lindholm *et al.* 2004. *Heredity* **92**, 156-162.
(3) Hurtado-Gonzales and Uy. 2010. *BMC Evol Biol* **10**, 391. (4) Liley. 1966. Behaviour Supplement **13**, 1-197.

**Table S2.** Sequencing method (10X Chromium linked reads or direct PE Illumina) and number of paired reads used for coverage analyses of all 40 wild caught individuals.

| Morph | Sample | Population | Sequencing | # Paired Reads |
|---|---|---|---|---|
| Females | P 15 | Cemetery | 10X | 566,474,152 |
| | P 26 | Seawall Trench | 10X | 586,091,672 |
| | P 32 | Cemetery | 10X | 642,872,944 |
| | P 34 | Seawall Trench | 10X | 599,429,008 |
| | P 36 | Cemetery | 10X | 587,807,726 |
| | P 37 | Seawall Trench | 10X | 586,019,978 |
| | P 16 | Cemetery | Illumina | 755,768,880 |
| | P 25 | Seawall Trench | Illumina | 861,217,302 |
| | P 33 | Cemetery | Illumina | 591,002,416 |
| | P 35 | Cemetery | Illumina | 813,848,774 |
| | P 39 | Seawall Trench | Illumina | 541,464,112 |
| Immaculata Morph | P 05 | Seawall Trench | 10X | 665,696,764 |
| | P 06 | Seawall Trench | 10X | 687,641,508 |
| | P 09 | Seawall Trench | 10X | 666,405,666 |
| | P 10 | Seawall Trench | 10X | 628,472,300 |
| | P 40 | Seawall Trench | Illumina | 492,568,288 |
| Parae Morph | P 03 | Seawall Trench | 10X | 636,937,278 |
| | P 04 | Seawall Trench | 10X | 673,606,290 |
| | P 27 | Seawall Trench | 10X | 682,154,596 |
| | P 29 | Seawall Trench | 10X | 697,480,156 |
| | P 30 | Seawall Trench | 10X | 598,808,486 |
| | P 31 | Seawall Trench | 10X | 634,574,988 |

| | | | | |
|---|---|---|---|---|
| | P 28 | Cemetery | Illumina | 799,757,386 |
| **Red Melanzona Morph** | P 01 | Cemetery | 10X | 605,264,908 |
| | P 02 | Cemetery | 10X | 599,899,768 |
| | P 11 | Seawall Trench | 10X | 663,454,532 |
| | P 12 | Cemetery | 10X | 679,401,488 |
| | P 19 | Cemetery | 10X | 619,323,522 |
| | P 20 | Seawall Trench | 10X | 614,834,422 |
| | P 21 | Cemetery | 10X | 564,829,682 |
| | P 38 | Seawall Trench | Illumina | 789,618,098 |
| **Yellow Melanzona Morph** | P 13 | Seawall Trench | 10X | 632,280,752 |
| | P 14 | West Watuka | 10X | 622,311,946 |
| | P 22 | Seawall Trench | 10X | 649,735,580 |
| | P 23 | West Watuka | 10X | 674,089,582 |
| | P 24 | West Watuka | 10X | 587,826,556 |
| **Blue Melanzona Morph** | P 17 | Cemetery | 10X | 705,952,894 |
| | P 07 | Seawall Trench | Illumina | 771,876,214 |
| | P 08 | Cemetery | Illumina | 479,310,754 |
| | P 18 | Seawall Trench | Illumina | 813,513,894 |

**Table S3.** Assembly statistics for each of the 29 genomes *de novo* assembled by Supernova. Bold samples indicate those used for follow-up coverage analyses. Genome size is expected to be ~730MB based on close relatives with known genome sizes (*Poecilia reticulata* 731Mb, *Xiphophorus helleri* 730Mb).

| Morph | Sample | Assembly Size (MB) | Scaffold N50 (kb) | No. Scaffolds ≥ 10kb | No. Scaffolds ≥ 1kb | Contig N50 (kb) | Phased N50 (kb) | Coverage |
|---|---|---|---|---|---|---|---|---|
| **Red Melanzona** | **P01** | **663.83** | **3201.95** | **2081** | **20376** | **38.36** | **2252.96** | **53X** |
| | P02 | 664.43 | 2150.00 | 2493 | 20516 | 38.78 | 1359.88 | 51X |
| | P11 | 562.45 | 44.07 | 17986 | 70941 | 23.36 | 70.98 | 52X |
| | P12 | 661.61 | 3683.18 | 2114 | 22269 | 37.97 | 2240.87 | 60X |
| | P19 | 582.84 | 47.86 | 17071 | 67134 | 23.08 | 100.55 | 48X |
| | P20 | 611.40 | 45.28 | 18734 | 63207 | 23.45 | 94.87 | 42X |
| | P21 | 665.11 | 1679.06 | 2677 | 20797 | 38.28 | 1266.47 | 51X |
| **Yellow Melanzona** | P13 | 654.30 | 1212.01 | 3205 | 23821 | 36.88 | 1120.80 | 53X |
| | P14 | 616.95 | 50.94 | 16711 | 58496 | 24.61 | 115.13 | 49X |
| | P22 | 665.85 | 61.08 | 15566 | 53875 | 27.00 | 137.91 | 46X |
| | P23 | 655.37 | 97.46 | 12443 | 43298 | 27.22 | 190.30 | 61X |
| | P24 | 234.10 | 17.09 | 13932 | 117930 | 13.32 | 5.82 | 45X |
| **Blue Melanzona** | P17 | 654.75 | 1995.50 | 2568 | 23989 | 33.84 | 1598.03 | 63X |
| **Parae Morph** | P03 | 655.50 | 93.43 | 12361 | 43771 | 28.68 | 165.34 | 51X |
| | **P04** | **658.87** | **599.09** | **4375** | **25416** | **34.50** | **599.94** | **55X** |
| | P27 | 667.40 | 274.95 | 6754 | 28937 | 30.94 | 312.62 | 59X |
| | P29 | 599.06 | 38.95 | 20291 | 68801 | 21.92 | 84.43 | 46X |
| | P30 | 341.91 | 20.87 | 17566 | 108816 | 15.66 | 40.56 | 39X |
| | P31 | 604.74 | 39.45 | 20427 | 65373 | 21.69 | 91.41 | 44X |
| **Immaculata** | P05 | 71.97 | 14.71 | 4740 | 138919 | 13.03 | 5.44 | 42X |
| | P06 | 623.35 | 46.83 | 18525 | 62105 | 24.07 | 105.78 | 53X |

| | | | | | | | | |
|---|---|---|---|---|---|---|---|---|
| | **P09** | **651.69** | **3308.41** | **2291** | **22613** | **35.38** | **2284.14** | **61X** |
| | P10 | 650.86 | 105.21 | 11446 | 39921 | 28.88 | 183.48 | 52X |
| **Female** | P15 | 69.89 | 15.53 | 4441 | 140510 | 13.68 | 3.91 | 44X |
| | P26 | 96.86 | 13.50 | 6848 | 144112 | 11.90 | 6.34 | 38X |
| | **P32** | **600.79** | **40.57** | **19565** | **62451** | **23.02** | **75.27** | **48X** |
| | P34 | 508.35 | 34.52 | 18503 | 79658 | 20.75 | 92.35 | 37X |
| | P36 | 329.51 | 23.33 | 15804 | 103659 | 16.40 | 35.28 | 18X |
| | P37 | 382.74 | 23.43 | 18121 | 95193 | 17.16 | 41.07 | 21X |

**Table S4.** Amount of sequence in scaffolds with >5 morph-mers. Bold indicates samples used for annotation.

| Morph | Sample | Number of Scaffolds | Amount of sequence |
|---|---|---|---|
| Melanzona Morph | **P01** | **77** | **995,370** |
| | P02 | 80 | 1,122,565 |
| | P11 | 85 | 865,095 |
| | P12 | 81 | 1,153,044 |
| | P13 | 92 | 1,610,202 |
| | P14 | 91 | 926,825 |
| | P17 | 104 | 4,250,903 |
| | P19 | 82 | 990,514 |
| | P20 | 94 | 877,954 |
| | P21 | 80 | 1,070,743 |
| | P22 | 95 | 1,164,800 |
| | P23 | 98 | 1,283,990 |
| | P24 | 113 | 668,430 |
| Parae Morph | P03 | 19 | 164,531 |
| | **P04** | **14** | **127,542** |
| | P27 | 14 | 84,778 |
| | P29 | 12 | 90,235 |
| | P30 | 9 | 96,497 |
| | P31 | 14 | 114,805 |
| Immaculata Morph | P05 | 126 | 521,712 |
| | P06 | 100 | 1,035,078 |
| | **P09** | **100** | **9,748,162** |
| | P10 | 81 | 1,388,117 |

**Table S5.** Genes annotated on the scaffolds with the Y-mers that are present in every male.

| Gene Name | Copies |
|---|---|
| CRYBB1 | 3 |
| CRYBA4 | 1 |
| GGTA1 | 1 |
| Retrovirus-related Pol polyprotein from type-2 retrotransposable element R2DM | |
| Retrovirus-related Pol polyprotein from type-1 retrotransposable element R2 | |

**Table S6.** *Melanzona morph*: Genes predicted on sample P01 (red melanzona) scaffolds containing >5 melanzona-mers and >1.5-fold melanzona male: female coverage. Scaffolds with >0.025X coverage by melanzona males but <0.025X coverage by females are considered 'Male Unique'. Scaffolds with >0.025X coverage by melanzona males but <0.025X coverage by non-melanzona males are considered 'Melanzona Unique'.

| Gene | P01 Scaffold | Melanzona:Female Coverage | Melanzona:Non-Melanzona Coverage |
|---|---|---|---|
| TRIM35 | 111589_P01 | Male Unique | Melanzona Unique |
| Tbx3 | 113310_P01 | Male Unique | 5.08 |
| Tbx3 | 113310_P01 | Male Unique | 5.08 |
| Texim2 | 114702_P01 | Male Unique | 14.57 |
| KAT7 | 115621_P01 | Male Unique | 4.09 |
| Retrovirus-related | 111503_P01 | Male Unique | 3.15 |
| Texim3 | 111503_P01 | Male Unique | 3.15 |
| Translation initiation factor IF-2 (low E) | 112167_P01 | 15.69 | 0.55 |
| Texim2 | 103684_P01 | 13.63 | 10.75 |
| Unknown | 103684_P01 | 13.63 | 10.75 |
| Unknown | 111891_P01 | 8.69 | 4.48 |
| LINE-1 type transposase | 115646_P01 | 3.78 | 3.52 |
| Texim2 | 112537_P01 | 1.67 | 2.12 |
| Unknown | 112537_P01 | 1.67 | 2.12 |
| Amyloid-beta A4 precursor | 116260_P01 | 1.91 | 0.97 |

**Table S7.** *Immaculata morph*: Genes predicted on sample P09 scaffolds containing >5 immaculata-mers and >1.5 fold immaculata male: female coverage. Scaffolds with >0.025X coverage by immaculata males but <0.025X coverage by females are considered 'Male Unique'. Scaffolds with >0.025X coverage by immaculata males but <0.025X coverage by non-immaculata males are considered 'Immaculata Unique'.

| Gene | P09 Scaffold | Immac:Female Coverage | Immac:Non-Immac Coverage |
|---|---|---|---|
| NLRC3 | 124510_P09 | Male Unique | Immaculata Unique |
| Trim39 | 125211_P09 | Male Unique | Immaculata Unique |
| Ty3 retrotransposon | 141715_P09 | Male Unique | 2.01 |
| MSI1 | 143420_P09 | Male Unique | 11.57 |
| Tbx3 | 143386_P09 | Male Unique | 2.40 |
| Trypsin-2 | 143386_P09 | Male Unique | 2.40 |
| R2DM retrovirus | 154564_P09 | Male Unique | 3.94 |
| Tbx3 | 140269_P09 | Male Unique | 1.62 |
| TBX3 | 87093_P09 | Male Unique | 0.94 |
| GPI-anchored protein 58 | 102953_P09 | 4.14 | 8.25 |
| NLRC3 | 139939_P09 | 4.06 | 7.46 |
| unknown | 139939_P09 | 4.06 | 7.46 |
| NLRC3 | 124079_P09 | 4.13 | 3.01 |
| Cetn3 | 137893_P09 | 1.57 | 1.15 |
| Retrovirus PABLB | 149075_P09 | 4.17 | 2.08 |
| Texim2 | 131979_P09 | 3.90 | 1.31 |
| unknown | 126759_P09 | 5.56 | 1.27 |
| unknown | 148405_P09 | 3.56 | 1.95 |

**Table S8.** Scaffolds containing >5 of the 59 Y-mers present in all males contain 8,547 transposable elements in their 30,558,901bp.

| Class/family | Copies identified by repeat masker |
|---|---|
| DNA | 264 |
| DNA/CMC-EnSpm | 297 |
| DNA/Crypton-V | 10 |
| DNA/Dada | 13 |
| DNA/Ginger-1 | 13 |
| DNA/IS3EU | 62 |
| DNA/Kolobok-T2 | 32 |
| DNA/MULE-MuDR | 27 |
| DNA/Maverick | 99 |
| DNA/Merlin | 31 |
| DNA/P | 5 |
| DNA/PIF-Harbinger | 63 |
| DNA/PIF-ISL2EU | 63 |
| DNA/PiggyBac | 12 |
| DNA/Sola-1 | 2 |
| DNA/TcMar | 8 |
| DNA/TcMar-ISRm11 | 34 |
| DNA/TcMar-Tc1 | 4626 |
| DNA/TcMar-Tc2 | 30 |
| DNA/TcMar-Tigger | 5 |
| DNA/Zisupton | 31 |
| DNA/hAT | 17 |
| DNA/hAT-Ac | 314 |
| DNA/hAT-Blackjack | 14 |
| DNA/hAT-Charlie | 225 |
| DNA/hAT-Tip100 | 75 |

| | |
|---|---|
| DNA/hAT-hAT5 | 35 |
| DNA/hAT-hAT6 | 11 |
| LINE/Dong-R4 | 120 |
| LINE/I | 54 |
| LINE/L1 | 77 |
| LINE/L1-Tx1 | 30 |
| LINE/L2 | 776 |
| LINE/Penelope | 14 |
| LINE/R2-Hero | 2 |
| LINE/RTE-BovB | 361 |
| LINE/Rex-Babar | 399 |
| LTR/Copia | 6 |
| LTR/ERV1 | 37 |
| LTR/Gypsy | 113 |
| LTR/Ngaro | 65 |
| LTR/Pao | 36 |
| RC/Helitron | 22 |
| Retroposon | 4 |
| SINE | 1 |
| SINE/tRNA-Core-RTE | 1 |
| SINE/tRNA-V | 7 |
| SINE/tRNA-V-RTE | 4 |

**Table S9.** Scaffolds containing >5 melanzona-mers contain 392 transposable elements in their 995,370bp.

| Class/family | Copies identified by repeat masker |
|---|---|
| DNA | 13 |
| DNA/CMC-EnSpm | 22 |
| DNA/Dada | 1 |
| DNA/IS3EU | 11 |
| DNA/Kolobok-T2 | 3 |
| DNA/Maverick | 10 |
| DNA/Merlin | 3 |
| DNA/PIF-Harbinger | 1 |
| DNA/PIF-ISL2EU | 1 |
| DNA/TcMar-Tc1 | 3 |
| DNA/Zisupton | 2 |
| DNA/hAT | 6 |
| DNA/hAT-Ac | 27 |
| DNA/hAT-Charlie | 28 |
| DNA/hAT-Tip100 | 2 |
| LINE/I | 2 |
| LINE/L1 | 24 |
| LINE/L1-Tx1 | 2 |
| LINE/L2 | 38 |
| LINE/RTE-BovB | 27 |
| LINE/Rex-Babar | 1 |
| LTR/Copia | 3 |
| LTR/ERV1 | 5 |
| LTR/Gypsy | 43 |
| LTR/Pao | 22 |
| RC/Helitron | 90 |
| SINE/tRNA-Core-RTE | 2 |

**Table S10.** Scaffolds containing >5 immaculata-mers contain 2,565 transposable elements in their 9,748,162bp.

| Class/Family | Copies identified by repeat masker |
|---|---|
| DNA | 103 |
| DNA/CMC-EnSpm | 104 |
| DNA/Crypton-H | 1 |
| DNA/Crypton-V | 7 |
| DNA/Dada | 4 |
| DNA/Ginger-1 | 3 |
| DNA/IS3EU | 47 |
| DNA/Kolobok-T2 | 13 |
| DNA/MULE-MuDR | 8 |
| DNA/MULE-NOF | 1 |
| DNA/Maverick | 25 |
| DNA/Merlin | 25 |
| DNA/P | 1 |
| DNA/PIF-Harbinger | 20 |
| DNA/Sola-1 | 1 |
| DNA/TcMar | 4 |
| DNA/TcMar-ISRm11 | 9 |
| DNA/TcMar-Tc1 | 1026 |
| DNA/TcMar-Tc2 | 21 |
| DNA/TcMar-Tigger | 4 |
| DNA/Zisupton | 8 |
| DNA/hAT | 12 |
| DNA/hAT-Ac | 145 |
| DNA/hAT-Blackjack | 2 |
| DNA/hAT-Charlie | 68 |
| DNA/hAT-Tip100 | 39 |
| DNA/hAT-hAT5 | 18 |
| DNA/hAT-hAT6 | 4 |
| DNA/hAT-hobo | 1 |
| LINE/Dong-R4 | 37 |
| LINE/I | 21 |
| LINE/L1 | 28 |
| LINE/L1-Tx1 | 7 |

| | |
|---|---|
| LINE/L2 | 340 |
| LINE/Penelope | 3 |
| LINE/R2-Hero | 1 |
| LINE/RTE-BovB | 112 |
| LINE/Rex-Babar | 112 |
| LTR/Copia | 1 |
| LTR/ERV1 | 12 |
| LTR/Gypsy | 55 |
| LTR/Ngaro | 14 |
| LTR/Pao | 9 |
| RC/Helitron | 38 |
| Retroposon | 1 |
| SINE/tRNA-Core-L2 | 45 |
| SINE/tRNA-V | 4 |
| SINE/tRNA-V-RTE | 1 |
